# VeTra: a tool for trajectory inference based on RNA velocity

**DOI:** 10.1101/2020.09.01.277095

**Authors:** Guangzheng Weng, Junil Kim, Kyoung Jae Won

**Affiliations:** Department of Biology, The bioinformatics Centre, University of Copenhagen, 2200 Copenhagen N, Denmark; Biotech Research and Innovation Centre (BRIC), University of Copenhagen, 2200 Copenhagen N, Denmark; Novo Nordisk Foundation Center for Stem Cell Biology, DanStem, Faculty of Health and Medical Sciences, University of Copenhagen, Ole Maaløes Vej 5, 2200 Copenhagen N, Denmark; Department of Bioinformatics, School of Systems Biomedical Science, Soongsil University, 369 Sangdo-Ro, Dongjak-Gu, 06978 Seoul, South Korea

**Keywords:** single cell RNA sequencing, Trajectory Inference, RNA velocity, Weakly Connected Component

## Abstract

Trajectory inference (TI) for single cell RNA sequencing (scRNAseq) data is a powerful approach to interpret dynamic cellular processes such as cell cycle and development. Still, however, accurate inference of trajectory is challenging. Recent development of RNA velocity provides an approach to visualize cell state transition without relying on prior knowledge. To perform TI and group cells based on RNA velocity we developed VeTra. By applying cosine similarity and merging weakly connected components, VeTra identifies cell groups from the direction of cell transition. Besides, VeTra suggests key regulators from the inferred trajectory. VeTra is a useful tool for TI and subsequent analysis.

## Introduction

Trajectory analysis using single cell transcriptomics is useful to understand temporal transition of cell states. By showing transcriptional changes along the obtained trajectory (or pseudo-time), TI has been applied to studying various biological processes including cell cycle, cancer development and cell differentiation (Tran & Bader, 2020).

Besides pseudo-temporal analysis, TI has been used to assist identifying gene regulatory rules. Visual inspection of gene expression changes along the inferred trajectory suggested potential regulators for the biological processes (Trapnell et al., 2014). Systemic approaches have been developed to reconstruct gene regulatory networks (GRNs) using the gene expression changes along the inferred trajectory (Kim, T. Jakobsen, Natarajan, & Won, 2021; Matsumoto et al., 2017). Well inferred trajectory helped improve the performance of GRN reconstruction (Qiu et al., 2020).

At present, more than 70 methods have been published to infer trajectory from scRNAseq data (Saelens, Cannoodt, Todorov, & Saeys, 2019). Majority of TI approaches have been developed based on the transcriptomic similarity among cells. Pseudo-time is obtained from the ordered cells based on the transcriptomic similarities (Saelens et al., 2019). However, transcriptomic similarity cannot specify the initial and the terminal points of the trajectory. When using TI algorithms based on the transcriptomic similarity, a user has to provide the direction of a cellular process, roots/terminals using the marker genes or any prior knowledge about experiments to obtain pseudo-time (Haghverdi, Büttner, Wolf, Buettner, & Theis, 2016; Setty et al., 2016; Trapnell et al., 2014).

It is more challenging to infer trajectory when cell dynamics are complex. It is difficult to assign an accurate fate (lineage) to a cell around branching regions. To improve robustness in detecting TI, there have been attempts to confine the inferred trajectory to a set of fixed topological structures, e.g. linear, bifurcation, and cycle (Saelens et al., 2019). However, the use of pre-defined structures can restrict the opportunity to identify new structures.

Recently, RNA velocity has been suggested to analyze scRNAseq data by incorporating mRNA dynamics (La Manno et al., 2018). Notably, RNA velocity can provide the direction and the speed of movement of individual cells and visualize the dynamic cell transitions without any prior knowledge. The direction information from RNA velocity provides a potential solution to determine the developmental trajectory of a cell around branching points. Therefore, RNA velocity can be useful in developing precise TI tools.

To obtain precise TI and subsequent analysis without prior information, we developed VeTra. VeTra identifies cell groups belonging to the same lineage and suggests potential regulators in the identified groups. Compared with previous undirected graph based methods such as minimum spanning tree (Street et al., 2018) and graph-partitioning algorithm (Wolf et al., 2019), VeTra builds directed graph from RNA velocity. VeTra identifies isolated cell transition paths by searching for weakly connected components (WCC) which represent the coarse-grained structure of connected community (An, Janssen, & Milios, 2004). The isolated paths are further grouped together using the hierarchical clustering algorithm to finally form potential development lineages. As the results, VeTra identifies cell transition trajectory and determines the memberships of cells to each trajectory even around branching points. VeTra further suggests key regulators along each trajectory by integrating the engine of TENET (Kim et al., 2021). VeTra is a user-friendly tool for TI and subsequent analysis.

## Results

### VeTra groups the cells belonging to the same stream of trajectory

VeTra is an RNA velocity based TI tool that enables accurate lineage tracing and subsequent analysis for gene regulation. VeTra performs lineage tracing from the root to the terminal states by grouping cells based on the similarity in direction of cell transition. This enables VeTra to perform TI without prior knowledge or pre-defined lineage topology.

VeTra reconstructs the pseudo-temporal order of cells based on the coordinates and the velocity vector of cells in the low-dimensional embedding. The velocity vectors are estimated by extrapolating the spliced/unspliced read ratio to the local neighboring cells (La Manno et al., 2018). Given velocity vectors (**Figure 1A**), VeTra reconstructs multiple directed graphs. To link cells based on transition, *k* nearest neighbours of a cell with similar direction are selected using cosine similarity (*cos*_*1*_) (**Figure 1B**, Methods). Among them, the nearby cell located upstream with the highest cosine similarity (*cos*_*2*_) is selected (**Figure 1B**, Methods). Once all cells were investigated for their next transition, multiple directed graphs are obtained (**Figure 1C**). To find a coarse-grained structure of the directed graph, VeTra identifies WCCs where every cell is reachable from every other cell regardless of the direction of relationships (Walker, 1992) (**Figure 1D**). The WCCs are grouped together when they are similar and close each other (Method) (**Figure 1E**). Finally, we obtained the pseudo-time ordering of cell groups (**Figure 1F**) by projecting the member cells onto the principal curve (Hastie & Stuetzle, 1989).

**Figure 1.**
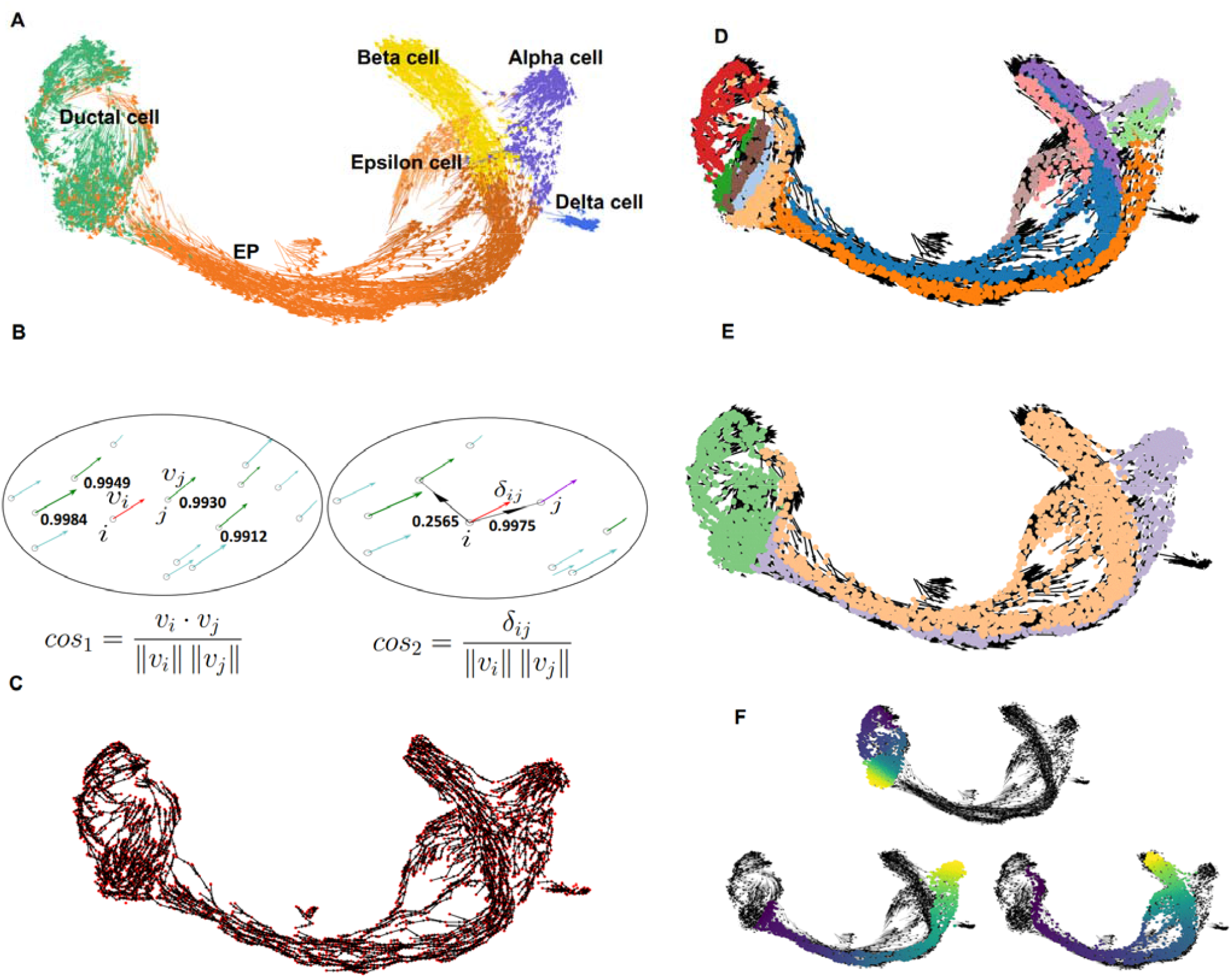
VeTra reconstructs single-cell trajectories for multiple cell lineages. **A**. A 2D embedding plot using the scRNAseq for pancreatic development. **B**. Cosine similarity to search for the neighboring cells with similar direction. *cos*_*1*_ finds the vectors with similar direction and *cos*_*2*_ identifies the cell to transit from a cell. **C**. The directed graph obtained by applying cosine similarity. **D**. The WCCs obtained using all possible paths. **E**. The grouped WCCs using a hierarchical clustering algorithm. **F**. The pseudo-time for each lineage identified by VeTra.

### VeTra’s trajectory matches well with the known lineage or biological process

We applied VeTra to infer the trajectory for various scRNAseq datasets with known cell dynamics structures for pancreatic development (Bastidas-Ponce et al., 2019; Bergen, Lange, Peidli, Wolf, & Theis, 2020), chromaffin cell differentiation (Furlan et al., 2017; La Manno et al., 2018), neural lineages in the hippocampus (Hochgerner, Zeisel, Lönnerberg, & Linnarsson, 2018; La Manno et al., 2018), and cell cycle (Xia, Fan, Emanuel, Hao, & Zhuang, 2019). For benchmarking we used Slingshot (Street et al., 2018), FateID (Herman & Grün, 2018) and PAGA (Wolf et al., 2019) since they showed best performance in recent benchmarking tests (Saelens et al., 2019). We also include CellRank (Bergen et al., 2020) and CellPath (Zhang & Zhang, 2021a) as they are developed based on RNA velocity. We used Diffusion pseudo-time (DPT) to provide pseudo-time ordering to CellRank. We provided known roots and terminals cell to run Slingshot and FateID.

#### Pancreatic development (a topology with crossing branches)

During the pancreatic development endocrine progenitors differentiate into alpha or beta cells (Bastidas-Ponce et al., 2019). Ductal cells are originated from common progenitors, which are fated to endocrine and exocrine lineages (Reichert & Rustgi, 2011). From the transcriptome of pancreatic development (E15.5), VeTra identified three major trajectory groups; i) alpha cell differentiation from endocrine progenitors (EP), ii) beta/epsilon differentiation from EP, and iii) ductal development (**Figure 2A**). The first two clusters were commonly originated from EPs and bifurcated into different lineages.

**Figure 2.**
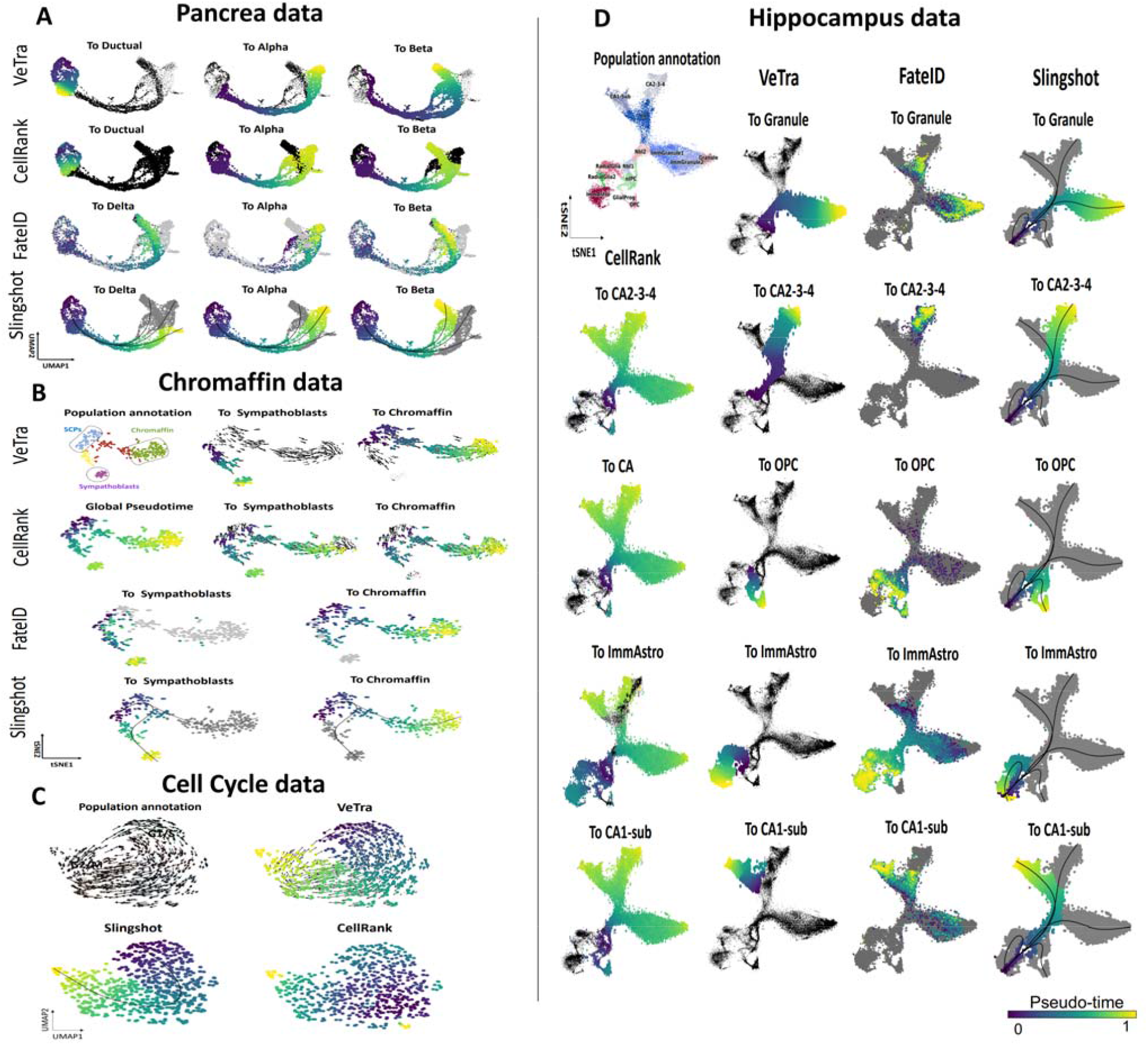
Performance assessment of the TI tools using the scRNAseq datasets with various topologies. The trajectories inferred by VeTra, CellRank, FateID and Slingshot using the scRNAseq data for **A**. pancreatic development. **B**. chromaffin and sympathoblasts development. **C**. cell cycle **D**. hippocampus development.

CellRank identified similar three trajectories with a slightly different starting point close to ductal cells for alpha and beta cell trajectories (**Figure 2A**). FateID identified the trajectory to alpha and beta cells but failed in identifying ductal lineage. Slingshot did not find delta/beta cell lineage as well as ductal cell lineage (**Figure 2A**). PAGA inferred coarse-grained trajectories from EP to alpha, beta, delta and epsilon cells but it also found trajectories between unrelated cell types (e.g. between alpha and beta cells) (Supplementary Figure 1). CellPath did not find the path to ductal cells (Supplementary Figure 2B).

#### Chromaffin cell development (a topology with diverging branches)

During the development, the chromaffin cells and sympathoblasts cells are generated from Schwann cell precursors (SCPs) (Furlan et al., 2017). Vetra, FateID and Slingshot successfully captured the branches towards sympathoblasts and chromaffin cells (**Figure 2B**). CellRank identified SCPs as initial states but failed to identify proper terminal states (**Figure 2B**). PAGA connected bridging cells presumably falsely as their fates are already committed to each branch (based on RNA velocity) (Supplementary Figure 1). CellPath only identified the path to the chromaffin cells (Supplementary Figure 2C).

#### Cell cycle (a cycling topology)

We applied TI tools on the scRNAseq dataset from U-2 OS cells (Xia et al., 2019) and investigate the topology for cell cycle. VeTra identified a single circular trajectory, starting at G1 phase towards G1/S, G2/M, and M phase, and finally going back to G1 phase (**Figure 2C**). Given the root cells in G1 phase, Slingshot identified similar trajectory. On the other hand, CellRank and CellPath estimated a linear-like trajectory (**Figure 2C**, Supplementary Figure 2D). PAGA connected cell populations with cycle, but inferred wrong trajectories between separated cell cycles (Supplementary Figure 1).

#### Hippocampus development (complex diverging branches)

We further evaluated the performance using the scRNAseq data for hippocampus development, which has been identified to have five diverging branches: astrocytes, oligodendrocyte progenitor cells (OPC), dentate gyrus granule neurons and pyramidal neurons including CA1, CA2, CA3 and subiculum (La Manno et al., 2018). Among these five main branches, astrocytes and OPCs are commonly branched from radial glial cells and the other neuronal branch cells are originated from immature neuroblast cells (La Manno et al., 2018).

VeTra identified five lineages towards CA1-subiculum, CA2-3-4, granule, astrocytes, OPC (**Figure 2D**). VeTra also predicted that CA2-3-4 cells and granules share the same neuroblastic origin (Nbl1) and astrocytes and oligodendrocyte progenitors share the same intermediate progenitor origin (nIPC), which is consistent with the original RNA velocity results (La Manno et al., 2018). CellRank identified one correct (Nbl1) and two incorrect (Granule, and CA1-Sub) initial states (**Figure 2D**). The five terminal states estimated by CellRank were CA, CA2-3-4, ImmAstro, GliaProg, and nIPC, including three potential misclassified intermediate states (GliaProg, nIPC and CA). Due to potential incorrect initial and terminal state assignment, the trajectory inferred by CellRank failed in distinguishing the five separated trajectories. Slingshot detected five lineages (**Figure 2D**) but determined wrong initial states (always from ImmAstro). FateID did not infer any meaningful trajectories (**Figure 2D**). PAGA inferred clear trajectories from Nbl1 to CA1-sub, CA2-3-4 and Granule. However, PAGA falsely connected ImmAstro and GlialProg (Supplementary Figure 1). CellPath showed severely truncated paths for this dataset (Supplementary Figure 2A).

### Performance evaluation using simulated datasets

The assessment using scRNAseq data demonstrated the outstanding performance of VeTra (**Figure 2**). However, the gold standard annotation for individual cells is usually not provided and it is hard to quantify the performance. For quantitative assessment, we simulated scRNAseq datasets using Dyngen (Cannoodt, Saelens, Deconinck, & Saeys, 2020) and generated three lineage structures: binary tree, trifurcation and converging. We additionally included a disconnected structure from VeloSim (Zhang & Zhang, 2021b). Beside visualization, we calculated the matching score by obtaining the average value of the normalized hamming distances (*D*) between the predicted and the known lineages. We used (1 − *D*) * 100 as the score.

VeTra successfully identified the identified lineages for four simulated datasets. Slingshot identified successfully except for the disconnected path. However, other approaches were not successful compared with VeTra or Slingshot (Supplementary Figure 3). The overall comparison (Supplementary Table 1) demonstrates the robustness of VeTra in TI.

**Figure 3.**
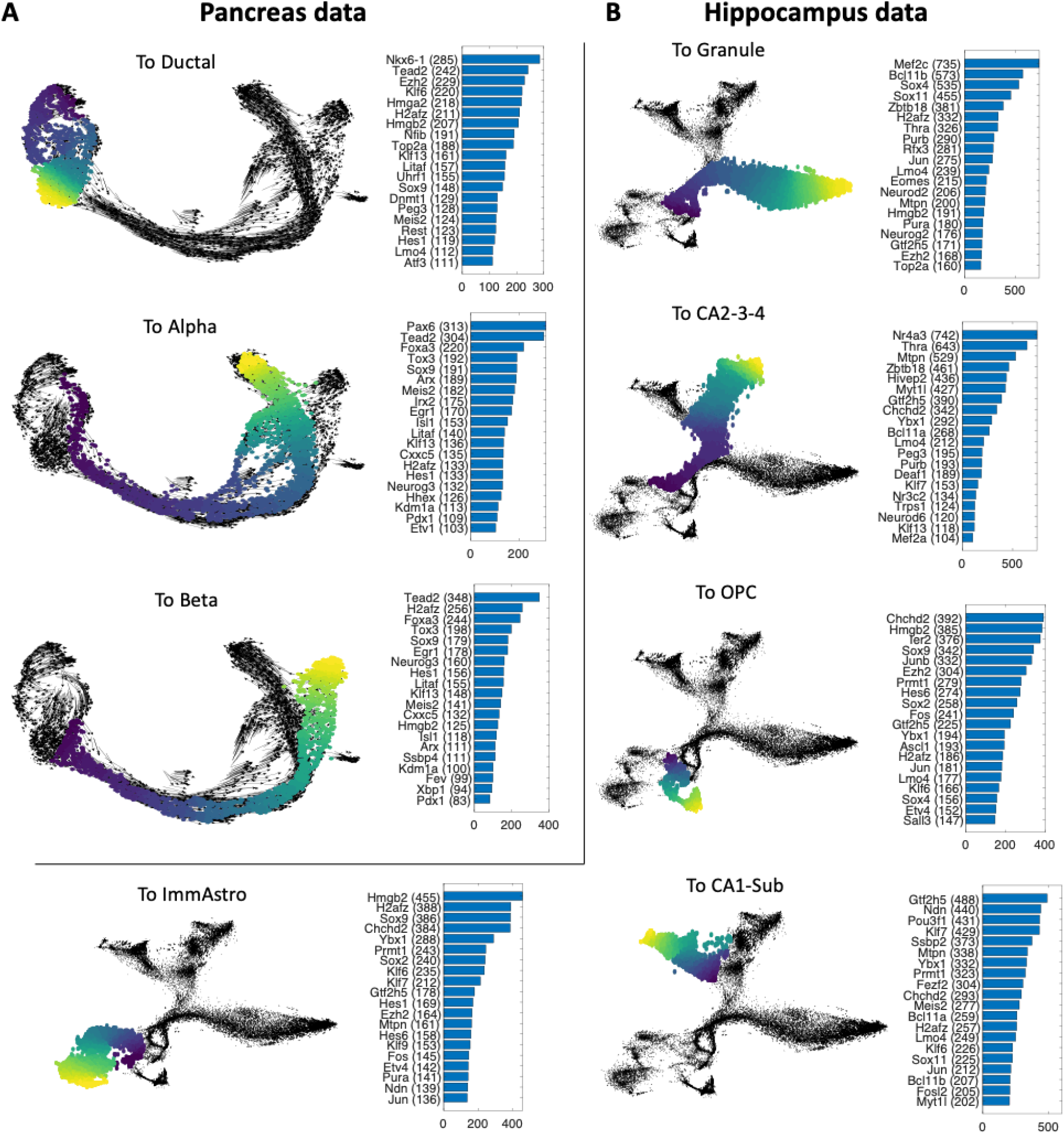
Key regulators for each trajectory identified by VeTra. **A**. The key regulators identified from the trajectories for ductal, alpha and beta cell development. **B**. The key regulators identified from the five developmental trajectories in mouse hippocampus.

### VeTra provides condition-specific master regulator

It has been studied that TI can influence the performance of GRN reconstruction. VeTra is equipped with a function to suggest key regulators by adopting the engine of TENET (Kim et al., 2021) that we previously developed to reconstruct GRN. TENET has a function to identify key regulators from the inferred causal relationships with their target genes (Kim et al., 2021). Incorporating TENET, VeTra has a function to identify most influential transcription factors (TFs) for the inferred trajectory.

Using the RNAseq dataset from pancreas development (Bastidas-Ponce et al., 2019), VeTra found key regulators for the three predicted major branches: alpha, beta and ductal (**Figure *3*A**). The key predicted regulators that distinguishing alpha cell development against others were Pax6, and Arx. Pax6 and Arx have been known to have a role during alpha cell development (Ashery-Padan et al., 2004; Gosmain, Cheyssac, Heddad Masson, Dibner, & Philippe, 2011). Pax6 regulates genes related to glucagon production (Ashery-Padan et al., 2004). Arx has been shown to play a similar role in alpha-cell development (Gosmain et al., 2011). VeTra also identified Neurog3 an endocrine (alpha and beta) specific key regulator (Gradwohl, Dierich, LeMeur, & Guillemot, 2000; Gu, Dubauskaite, & Melton, 2002; Schwitzgebel et al., 2000).

In addition, VeTra revealed condition-specific regulators for the five lineages of hippocampus development (**Figure *3*B**). For instance, Nr4a3 (Nor-1) is predicted a CA2-3-4 specific regulator by VeTra. Nor-1 deficient mice appear to have abnormal pyramidal cell layer in CA1 to CA3 (Pönniö & Conneely, 2004).

## Methods

To pick the most appropriate cell to which the vector of the cell *i* is pointing, *k* closest neighbor cells are collected from the head of the vector of a cell *i* in the low-dimensional space (**Figure 1B**). Among the neighbor cells, cells with similar direction are selected using a cosine similarity criterion between the cell *i* and the neighbor cell *j* 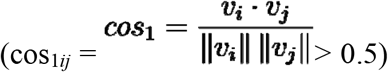 (**Figure 1B**), where *v*_*i*_ and *v*_*j*_ denote velocity vectors (2D coordinates) of cell *i* and cell *j*. The cell with the highest similarity for cos_2*ij*_ (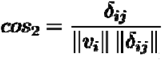, where δ_ij_ is the vector from *v*_*i*_ to *v*_*j*_) located upstream of the cell *i* is finally selected to obtain a directed graph in the same stream of trajectory (**Figure 1B**).

The hierarchical clustering is applied to the WCCs for further grouping (**Figure 1E**). The distance between the two sub-graphs is defined by the maximum distance of all the closest pairs of cells. To calculate the distance between cells, we calculated the Euclidean distance in the four-dimensional space (two dimensions using the gene expression and the other two using the velocity vector in the reduced dimensional space). To obtain full trajectory (from the root to the branch), we extended the memberships if a cell located nearby is similar (cos_1*ij*_ > 0.7). We obtained the pseudo-time ordering by projecting the member cells onto the principal curve (Hastie & Stuetzle, 1989).

## Discussion

TI is a widely used approach to understand temporal dynamics of cells from scRNAseq data. A number of approaches have already been developed to infer the trajectories. As shown in the examples using the simulated as well as real scRNAseq data (Figure 2 and Supplementary Figure S1-3), however, there is still room for algorithmic improvement for accurate detection of trajectory. We have noticed that the results of current TI tools do not always match with the cell dynamics observed by RNA velocity (La Manno et al., 2018). RNA velocity, by providing the direction of cell transition, enabled visual inspection of cell trajectory. VeTra is designed to group cells in accordance with the cell transition drawn by RNA velocity. To fully use the vector space provided by RNA velocity, VeTra used cosine similarity and obtained cell groups. The use of cosine similarity makes VeTra rely on the vector space for TI.

Our benchmarking test on the scRNAseq datasets exemplifies that VeTra successfully identified the trajectory drawn by RNA velocity and produced reasonable results that matched with the known topological structures for cell development and biological processes. It is interesting that VeTra outperformed other RNA velocity based methods such as CellRank (Bergen et al., 2020) and CellPath (Zhang & Zhang, 2021a) in our tests. Instead of using RNA velocity directly, CellRank calculates the probabilities of each cell transition to all terminals from initials. Therefore, the terminal states are not deterministic for CellRank. This may have influenced the performance of CellRank. CellPath uses meta-cells to obtain smoothed RNA velocity and reduce the computing cost (Zhang & Zhang, 2021a). In our test, CellPath has failed to annotate large number of cells for each lineage. A meta-cell is a cluster of cells. We suspect that meta-cells may smooth cell direction too much when cells exhibit diverse direction. Compared with them, VeTra tried to follow the velocity vector using cosine similarity without further processing them.

As the results, VeTra relies on RNA velocity calculation. RNA velocity can heavily depend on the correct measurement of spliced and unspliced RNA abundance. A recent study showed that the choice of experimental setting can change the results of RNA velocity (Soneson, Srivastava, Patro, & Stadler, 2021). As a downstream analysis tool of RNA velocity, VeTra can use the output from Velocyto (La Manno et al., 2018) or scVelo (Bergen et al., 2020). A previous study shows that the results of Velocyto and scVelo can be different (Cannoodt et al., 2020), which can also cause different results for VeTra. We investigated VeTra’s output after running Velocyto and scVelo. For the cell cycle and chromaffin development datasets, both Velocyto and scVelo showed similar results. However, we found different results for the hippocampus and the pancreas development datasets (Supplementary Figure 4). For VeTra, CellRank and CellPath, we selected the RNA velocity approach that produces the better performance for each dataset.

VeTra is equipped with the tool to predict key regulators by adopting the gear of TENET, a GRN reconstructor based on TE. It was discussed previously that TENET is capable of detecting key regulators using the aligned scRNAseq (Kim et al., 2021). VeTra equipped with the TENET engine will be useful for downstream analysis of TI.

VeTra requires the 2D coordinates embedded from expression profile and the RNA velocity vectors for each cell. A user can specify the number of groups. The output files include i) 2D embedding figures colored by grouping of cells, ii) 2D embedding figures colored by pseudo-time ordering, iii) lists of selected cells for each group, and iv) the pseudo-time ordering information for each group.

## Availability

Vetra is available at https://github.com/wgzgithub/VeTra.

## Funding

The Novo Nordisk Foundation Center for Stem Cell Biology is supported by a Novo Nordisk Foundation grant No. NNF17CC0027852. KJW is supported by the Lundbeck Foundation’s Ascending Investigator grant R313-2019-421 and the Independent Research Fund Denmark (0135-00243B).

## Competing interests

The authors declare no competing interests.

## Notes

### Competing Interest Statement

The authors have declared no competing interest.

