## Supplementary material for "VeTra: a tool for trajectory inference based on RNA velocity"

### VeTra: a tool to group cells based on dynamic trajectory

### Supplementary Materials

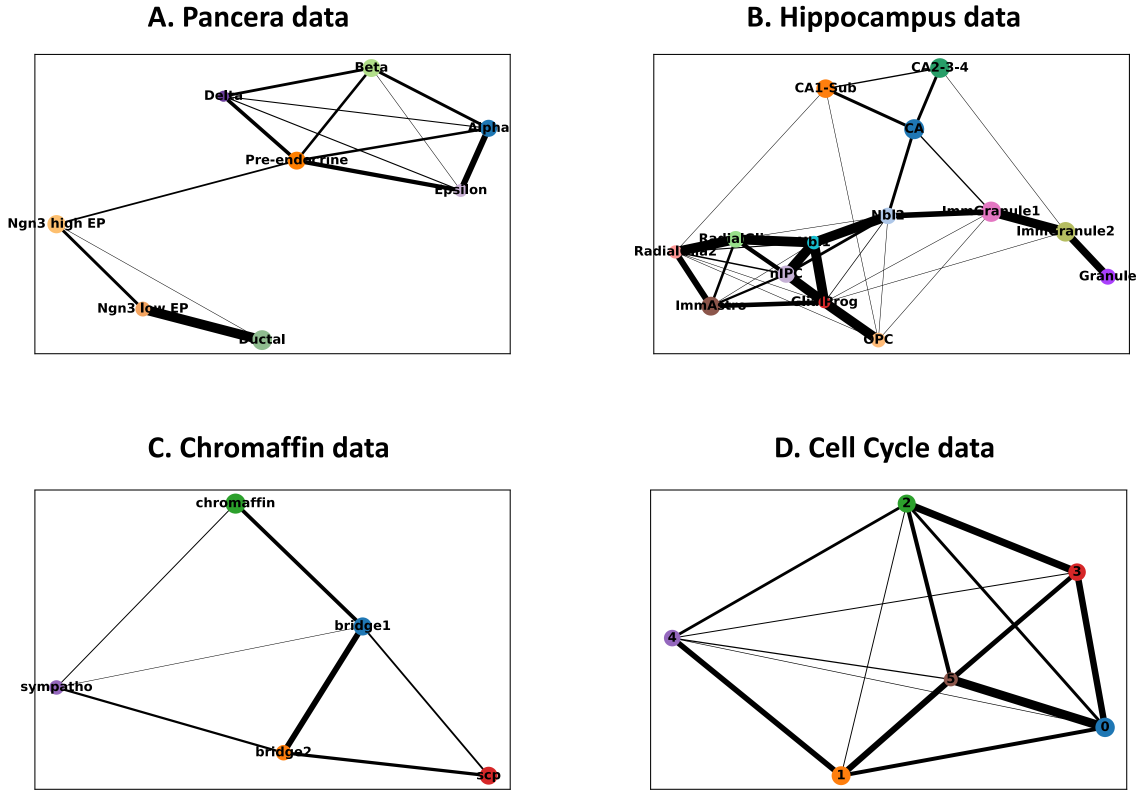


**Supplementary Figure 1.** The TI results from PAGA for various scRNAseq datasets

### Cellpath performance on real datasets

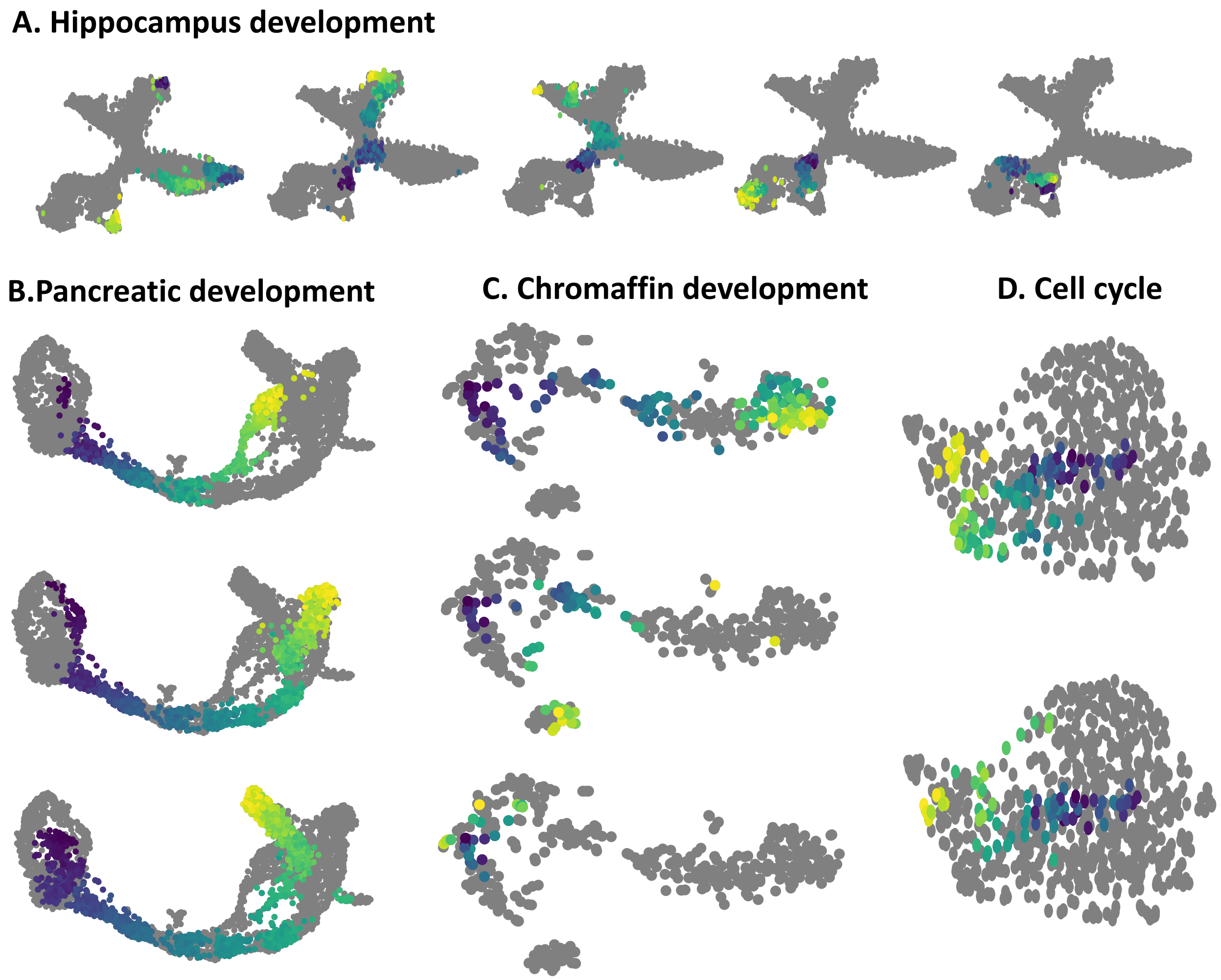


**Supplementary Figure 2.** The TI results from CellPath for various scRNAseq datasets

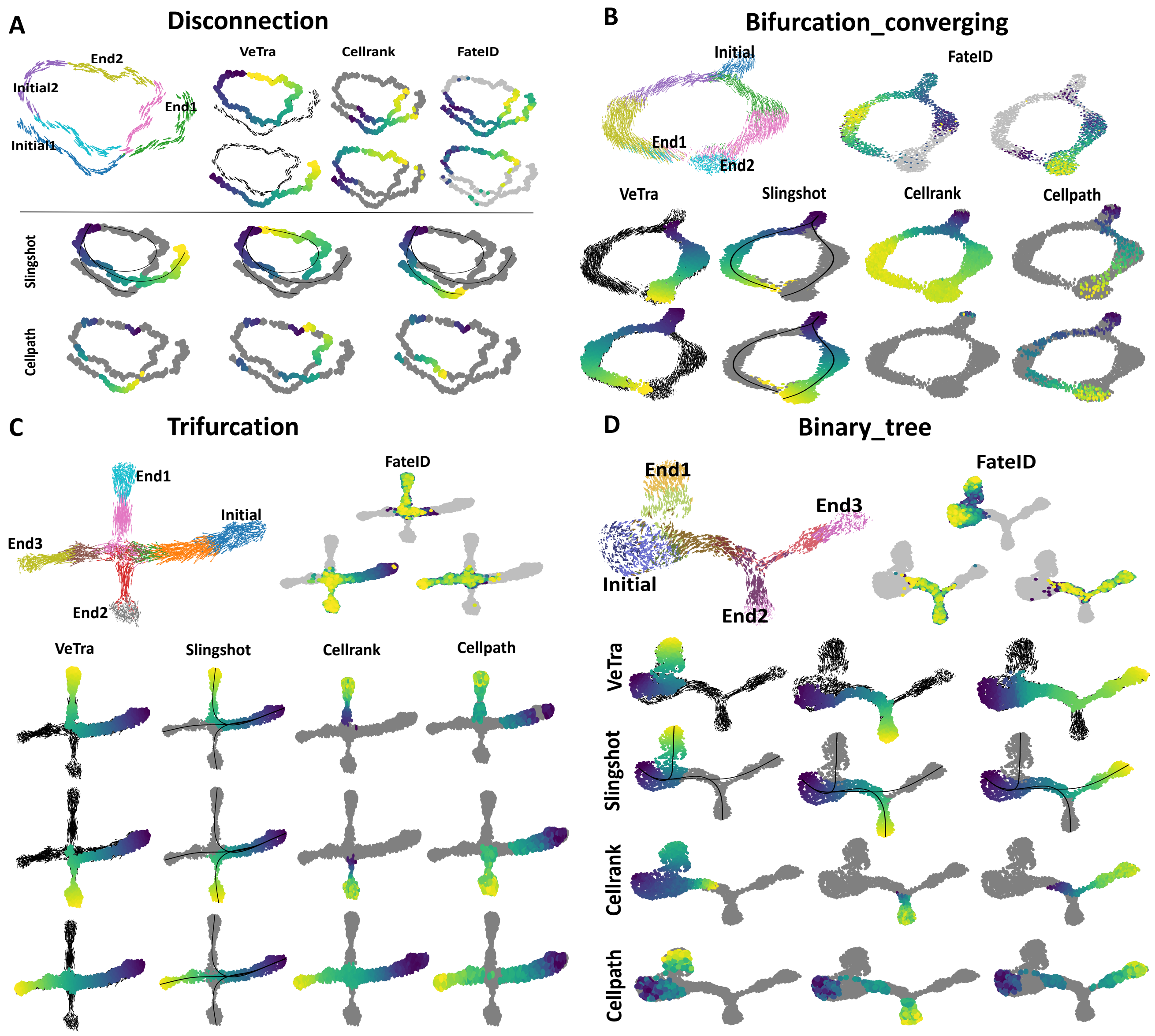


**Supplementary Figure 3.** Performance of all TI tools on the 4 types of simulated datasets

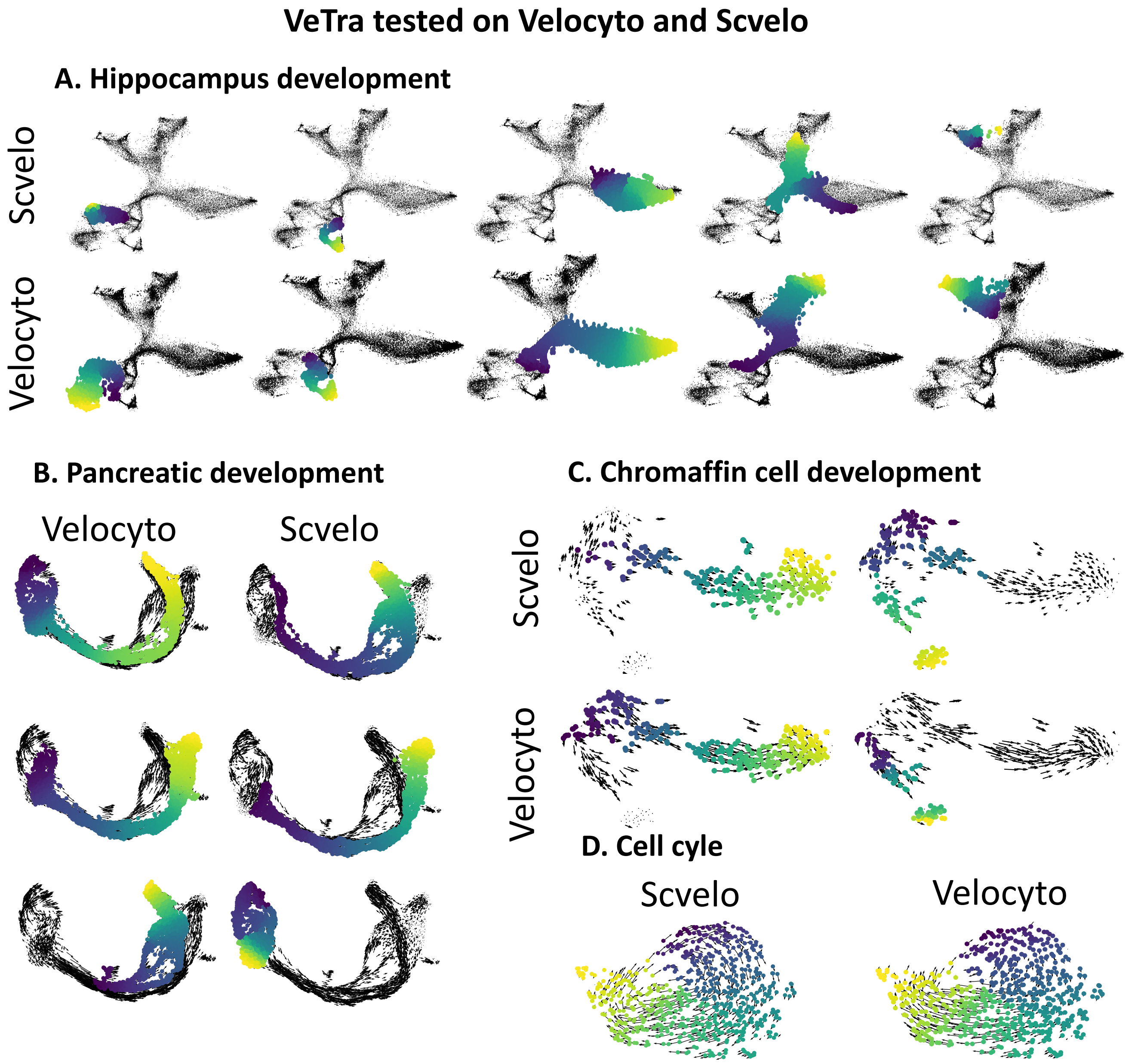


**Supplementary Figure 4.** VeTra's outputs tested on Velocyto and Scvelo

**Supplementary Table 1**. Performance assessment using the simulated datasets. The score is $(1 - D) * 100$, where $D$ is the average value of hamming distances between predicted lineages and the known lineages normalized to the total number of cells.

| | Trifurcation | Converging | Binary Tree | Disconnection | Overall |
|---|---|---|---|---|---|
| VeTra | **91** | **88** | **91** | **99** | **92** |
| Slingshot | **95** | **95** | **95** | **78** | **91** |
| FateID | **72** | **77** | **76** | **67** | **73** |
| CellRank | **71** | **52** | **69** | **63** | **64** |
| Cellpath | **78** | **56** | **64** | **67** | **66** |
